# Survival of Captive Chimpanzees: Impact of Location and Transfers

**DOI:** 10.1101/068858

**Authors:** Michael S. Lauer

## Abstract

To inform the retirement of NIH-owned chimpanzees, we analyzed the outcomes of 764 NIH-owned chimpanzees that were located at various points in time in at least one of 4 specific locations. All chimpanzees considered were alive and at least 10 years of age on January 1, 2005; transfers to a federal sanctuary began a few months later. During a median follow-up of just over 7 years, there were 314 deaths. In a Cox proportional hazards model that accounted for age, sex, and location (which was treated as a time-dependent covariate), age and sex were strong predictors of mortality, but location was only marginally predictive. Among 273 chimpanzees who were transferred to the federal sanctuary, we found no material increased risk in mortality in the first 30 days after arrival. During a median follow-up at the sanctuary of 3.5 years, age was strongly predictive of mortality, but other variables – sex, season of arrival, and ambient temperature on the day of arrival – were not predictive. We confirmed our regression findings using random survival forests. In summary, in a large cohort of captive chimpanzees, we find no evidence of materially important associations of location of residence or recent transfer with premature mortality.

## Introduction

Scientists have chosen to study chimpanzees to research a variety of topics including hepatitis, retrovirus immunodeficiency infection, brain function and development, and comparative genomics. NIH-funded researchers have relied historically on a small number of primate facilities as sources of chimpanzees in the United States. These include the Michale E. Keeling Center for Comparative Medicine and Research (Bastrop, Texas), the Texas Biomedical Research Institute/Southwest National Primate Research Center (San Antonio, Texas), and the Alamogordo Primate Facility, which is a research reserve facility. Animals no longer used in research have been housed in the federal chimpanzee retirement sanctuary (managed by Chimp Haven, Inc. in Keithville, Louisiana). Some NIH-funded studies have historically relied on captive chimpanzee populations that reside in other locations, including zoos, academic institutions, or other privately owned facilities.

In June 2013, the NIH decided to substantially reduce the use of chimpanzees in research.^1^ The NIH instituted a number of procedures and processes to phase out ongoing research and to retire all but 50 chimpanzees which would be retained for emerging research needs. In November, 2015, the NIH, upon further evaluation, decided to retire all of its chimpanzees and to disallow all invasive research.^2^

The NIH is actively working to retire all NIH-owned and -supported chimpanzees to the federal sanctuary at Chimp Haven. The retirement requires physical relocation of animals from the facilities where they are currently housed. The NIH chimpanzee population is aging and has an increasing incidence of various chronic diseases. To inform our retirement plan, we analyzed available outcomes data on NIH-owned animals from the three NIH-supported facilities and the federal sanctuary.

## Methods

### Sample, exposure, and outcomes

We followed the outcomes of NIH-owned captive chimpanzees who were alive and at least 10-years old on January 1, 2005. We chose this date as “time zero” because transfers to the federal sanctuary (Chimp Haven) began several months later and because NIH did not own any chimpanzees in 2005 that were less than 10 years of age. For each chimpanzee we had data available to us on date of birth, sex, locations at specific points in time, and, when applicable, date of death. The primary outcome was all-cause mortality as a function of age, sex, and location, with location considered as a time-dependent covariate.

### Analyses

For the primary outcome, we focused on animal-location intervals that were each bounded by arrival and end times. The arrival time (or “time 1”) was January 1, 2005 or the date of arrival to the location, whichever was later. The end time (or “time 2”) was the date of death, date of transfer to another location, or, as a right-censoring value, July 15, 2016 if the animal was still alive. Because location is a time-dependent covariate, some animals contribute “animal-time” information to more than one location.

We used two methods to assess the associations of age, sex, and location (as a time-dependent covariate) with all-cause mortality. We constructed a Cox proportional hazards model for each animal-location interval:

*S* (time 1, time 2, outcome_(Death=1, Censored=0)_) ~α_1_^*^age + β_1_^*^sex_(Male=1, Female=0)_ +χ_1_^*^location

Given the number of variables considered (age, sex, and 4 possible locations), we chose a P-value of 0.01 to declare statistical significance. For this model, we used the calculated age on January 1, 2005. We tested for possible non-linear associations using restricted cubic splines and we tested for possible interactions between covariates. We verified the Cox proportional hazards assumption by calculating scaled Schoenfeld residuals.

We also used random survival forests to assess the associations of age, sex, and location with all-cause mortality. Random survival forests, as we have described previously^3-6^, are an extension of Breiman’s random forests, but applied to right-censored data. Random forests allow for robust assessment of variable importance using machine-learning methods, which function independent of parametric assumptions and which account intrinsically for interactions, correlations, and outliers. We constructed 1000 trees (bootstrap samples) for each forest; for the age covariate, we calculated age on arrival.

In a secondary analysis, we analyzed all-cause mortality at Chimp Haven according to age on arrival, sex, season of arrival, and the high- and low-ambient temperatures in the Chimp Haven area on the day of arrival. We defined four seasons as: Winter – January, February, and March; Spring – April, May, and June; Summer – July, August, and September; Fall – October, November, and December. We obtained date-specific values for high and low temperature at Keithville LA from Weather Underground’s historical archive.7 We constructed Kaplan-Meier survival plots as well as parametric hazard function plots to assess mortality risk over time since arrival. We assessed the importance of candidate predictive variables with standard Cox proportional-hazards models and by random survival forest regressions.

We conducted our analyses with R 3.3.0, using the packages rms, Hmisc, ggplot2, dplyr, GGally, survival, and randomForestSRC.

## Results

We obtained data on 764 unique chimpanzees (366 or 48% male, 398 or 52% female). On January 1, 2005, the median age was 22.63 years (interquartile range 17.19 to 30.14; full range 10.09 to 51.06). During a median follow-up of 7.14 years, there were 314 deaths.

Table 1 shows characteristics according to animal-location intervals. Animals at Bastrop and Chimp Haven were somewhat older. As expected, times of follow-up at Chimp Haven were shorter since all animals there were transferred from other locations after January 1, 2005 (median time after January 1, 2005 for transfer 3.1 years, interquartile range 1.8 to 9.1 years).

**Table 1.** Characteristics according to animal-location intervals

| **Characteristic** | **APF**<br>**N=253** | **Bastrop**<br>**N=145** | **CH**<br>**N=273** | **SNPC**<br>**N=222** |
| --- | --- | --- | --- | --- |
| **Age** | 22 (18-29) | 24 (17-35) | 24 (17-35) | 22 (16-26) |
| **Sex (M)** | 54% (137) | 44% (64) | 45% (124) | 43% (96) |
| **Follow-up time** | 11.5 (5.5-11.5) | 7.7(5.6-9.2) | 3.5 (2.2-8.4) | 7.8 (2.3 – 11.5) |
| **Death** | 33% (83) | 39% (57) | 39% (107) | 30% (67) |

Continuous variables shown as median (25<sup>th</sup> percentile – 75<sup>th</sup> percentile).
APF = Alamogordo Primate Facility; CH = Chimp Haven; SNPC = Southwest National Primate Research Center

Table 2 shows the results of Cox proportional hazards modeling. All associations were log-linear and there were no significant interactions. The strongest predictor, by far, of mortality was age (as calculated to be on January 1, 2005), followed by male sex and location. Older age predicted higher mortality (adjusted hazard ratio comparing animals 30 years versus 17 years 2.23, 95% CI 1.91 to 2.61); males also had higher mortality (adjusted hazard ratio 1.50, 95% CI 1.20 to 1.88). Location was only marginally associated with mortality (Wald χ^2^=10, df=3, P=0.017). Compared to Chimp Haven, mortality was lower at APF (adjusted hazard ratio 0.65, 95% CI 0.48 to 0.88), while it was similar at Bastrop (adjusted hazard ratio 0.84, 95% CI 0.60 to 1.16) and almost identical at SNPC (adjusted hazard ratio 1.00, 95% CI 0.71 to 1.39).

**Table 2.**
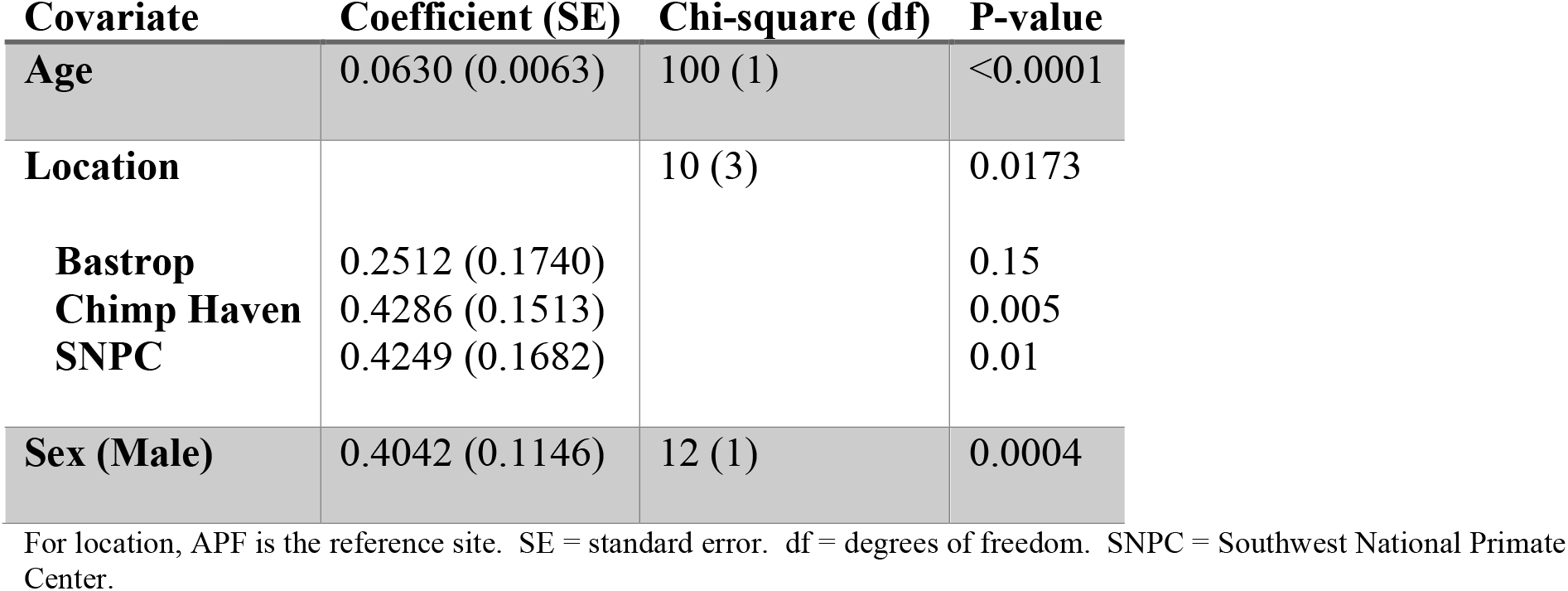
Cox proportional hazards model of mortality according to age, sex, and location (as a time-dependent covariate).

Figure 1 shows the results of modeling by random survival forests; the regression survival model had a c-statistic equivalent of 0.62. Consistent with the Cox proportional hazards model, age was by far the most important predictor of mortality, while location was the least important predictor.

**Figure 1.**
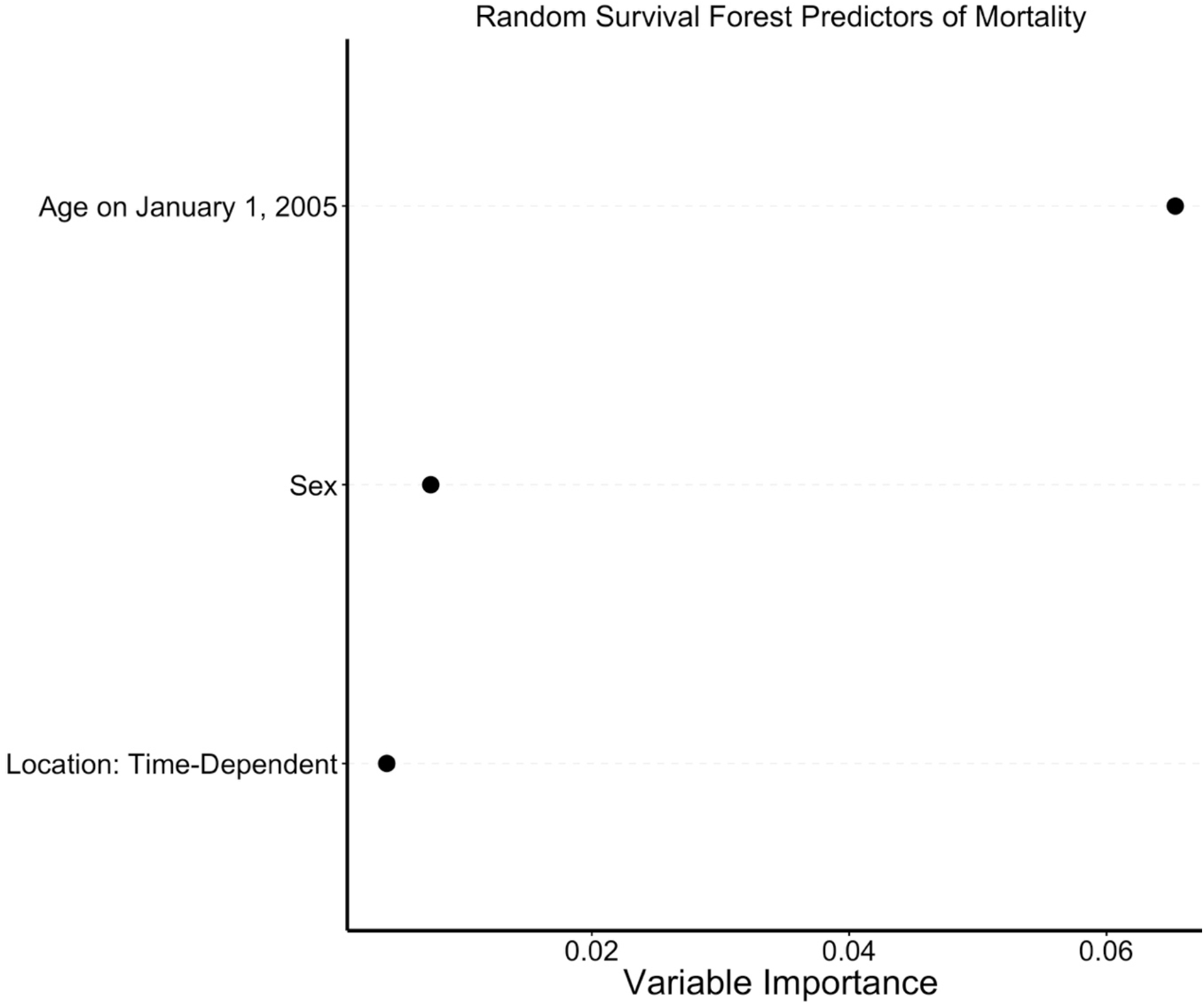
Results of Random Survival Forest regression. The plot shows variable importance (“VIMP”) values for each covariate.

Among the 273 chimpanzees who were at some point located at Chimp Haven, there were 107 deaths. Figure 2a shows the month of arrival, while Figures 2b and 2c show season and high-temperature values. More than half of transfers occurred in the months of April and May, while summer transfers were unusual.

**Figure 2.**
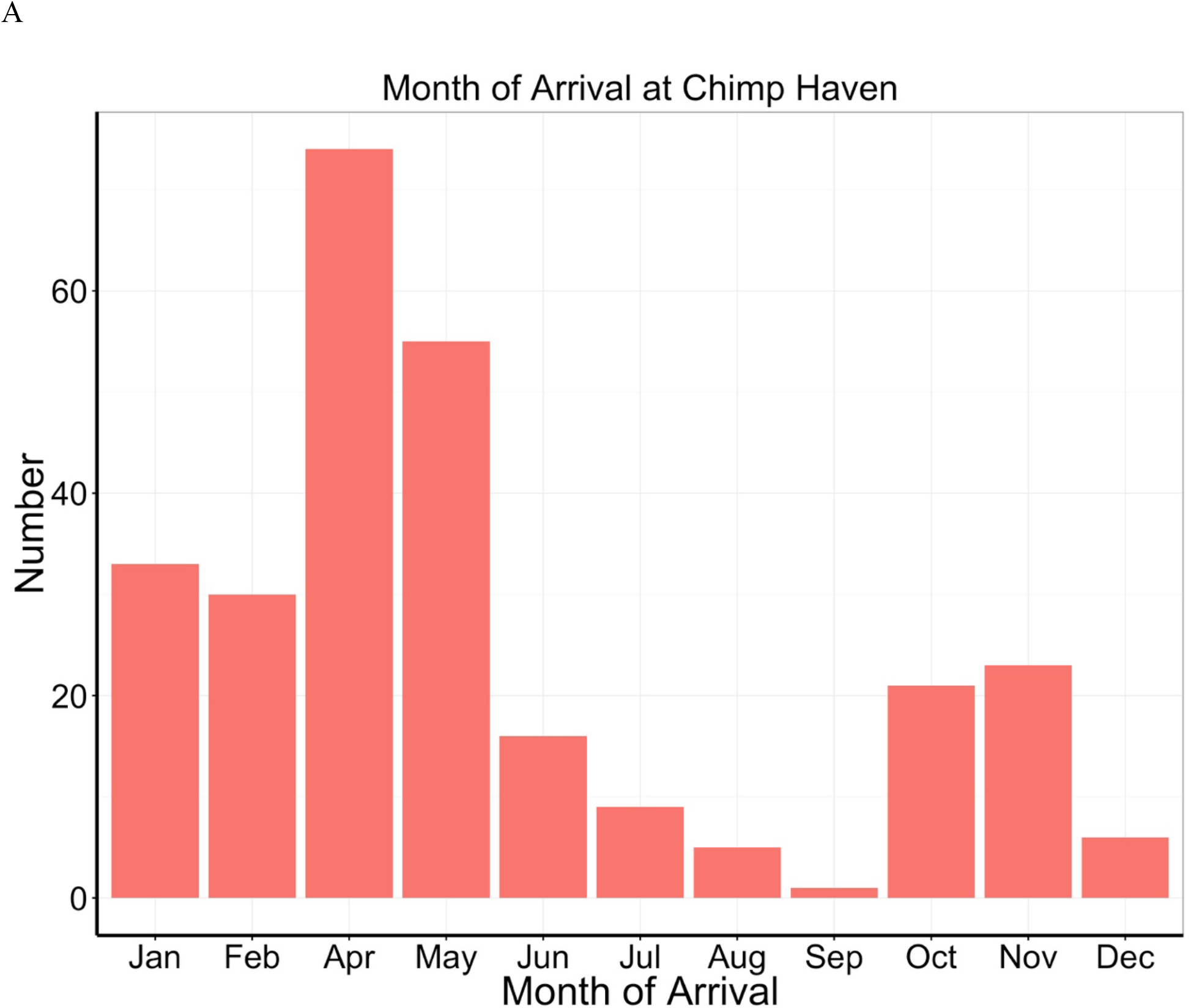
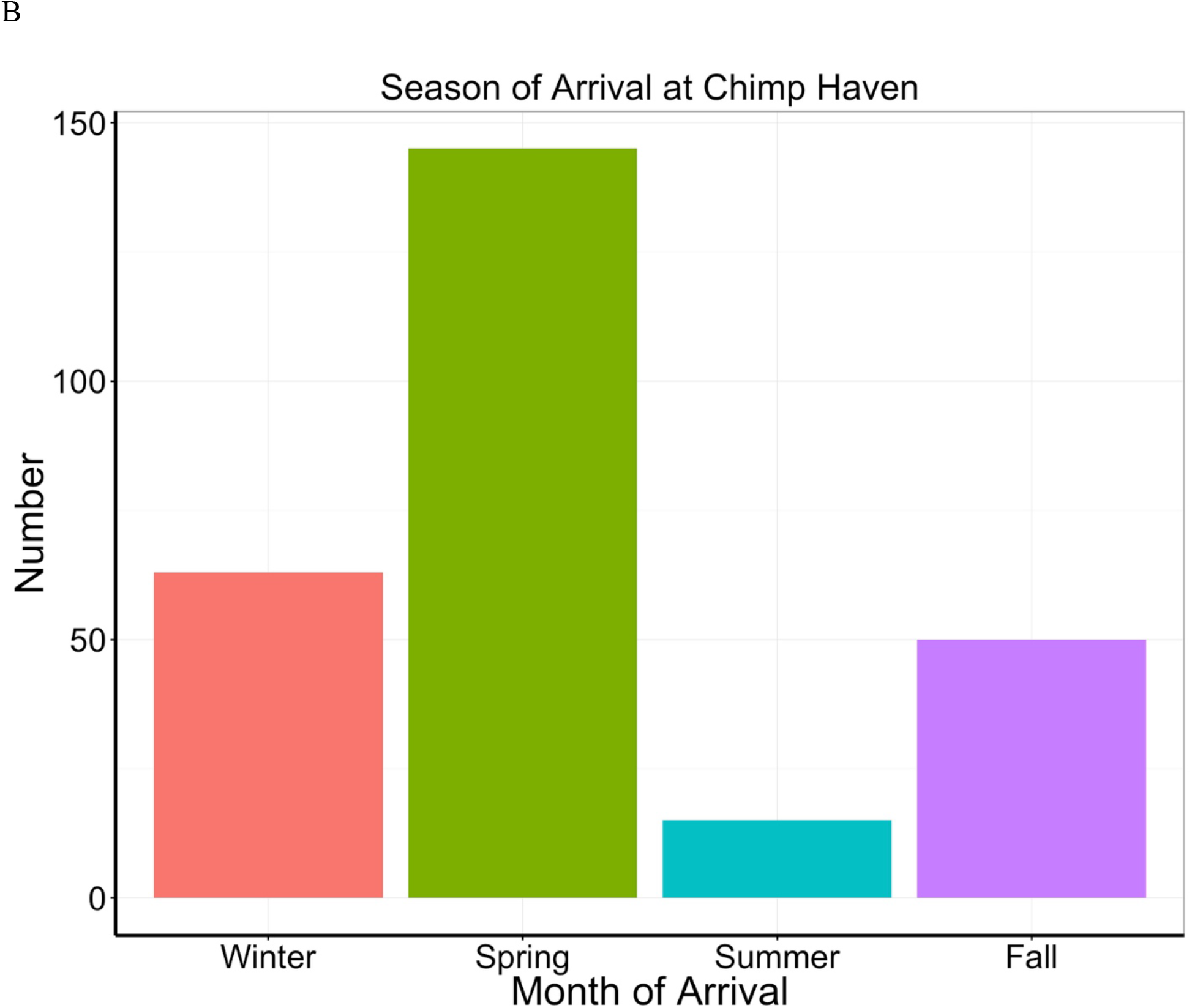
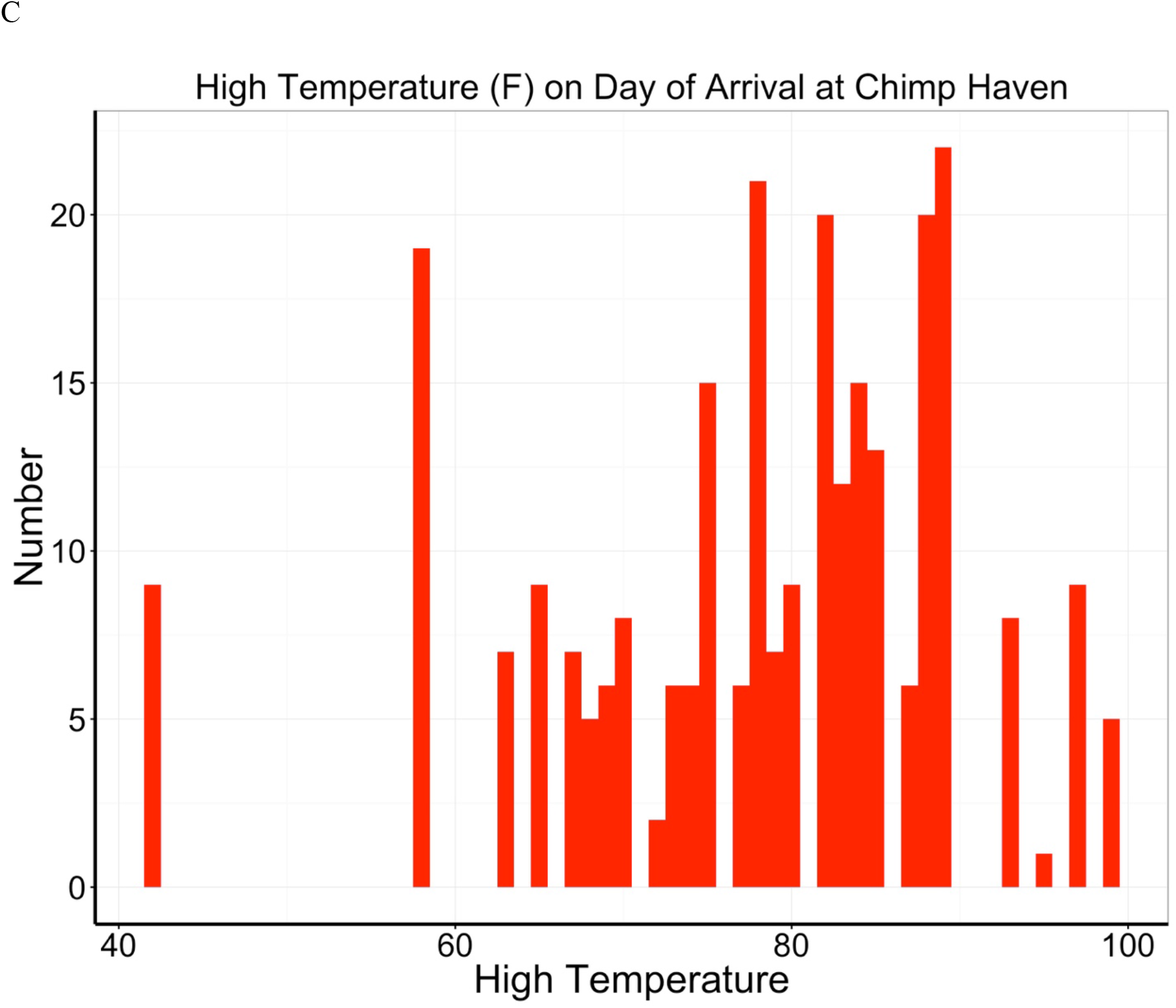
Time and conditions of arrival for 273 chimpanzees transferred to Chimp Haven. Panels A and B are histogram based on month and season of arrival. Panel C shows the high-temperature (in degrees Fahrenheit) recorded on the day of arrival.

In a Cox proportional hazards model, the only significant predictor of mortality at Chimp Haven was age at arrival (β=0.0679, se=0.0109, Wald χ^2^=39, P<0.0001). Other predictors were not significantly associated with time to death, including sex (P;0.16), season (0.88), or, in a separate model, high temperature on arrival (P=0.12). Figure 3 shows the results of a random survival forest regression; again age was by far the strongest predictor of mortality.

**Figure 3.**
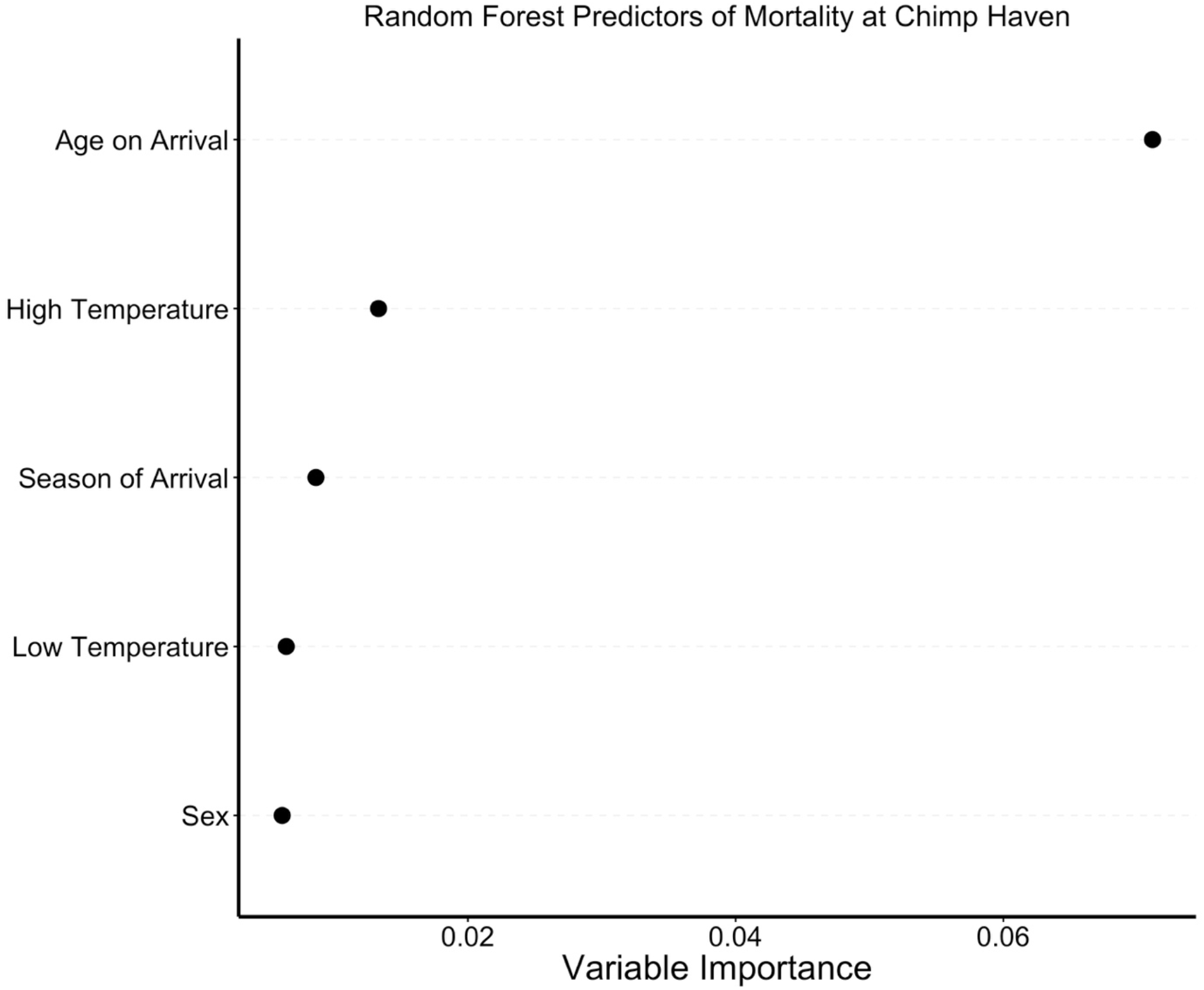
Results of Random Survival Forest regression for 273 chimpanzees transferred to Chimp Haven. The plot shows variable importance (“VIMP”) values for each covariate. “Arrival” refers to arrival date to Chimp Haven, which was time zero for these analyses.

Figure 4a shows survival, as measured by the Kaplan-Meier method, since arrival at Chimp Haven. There appears to be a monotonic pattern with no clear increased mortality risk early after arrival. Figure 4b focuses on the first year after arrival; there is a slightly greater decline in survival early on (within 30 days) reflecting the deaths of 3 animals, who were ages 30, 42, and 46 on arrival. Over up to 11 years of follow-up time, hazard for mortality increased from ~5% per year to 10% per year, as expected with the aging of the animals.

**Figure 4.**
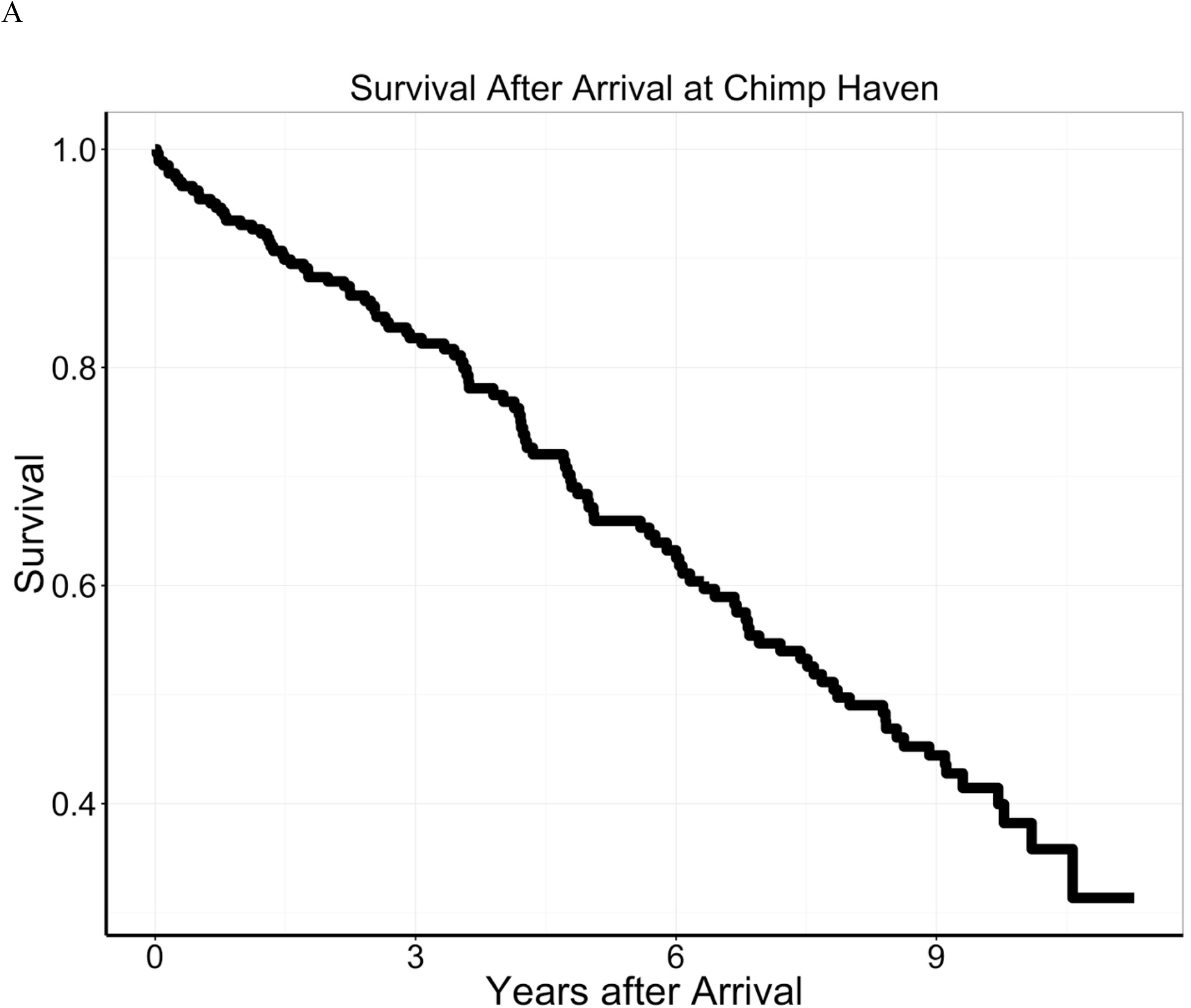
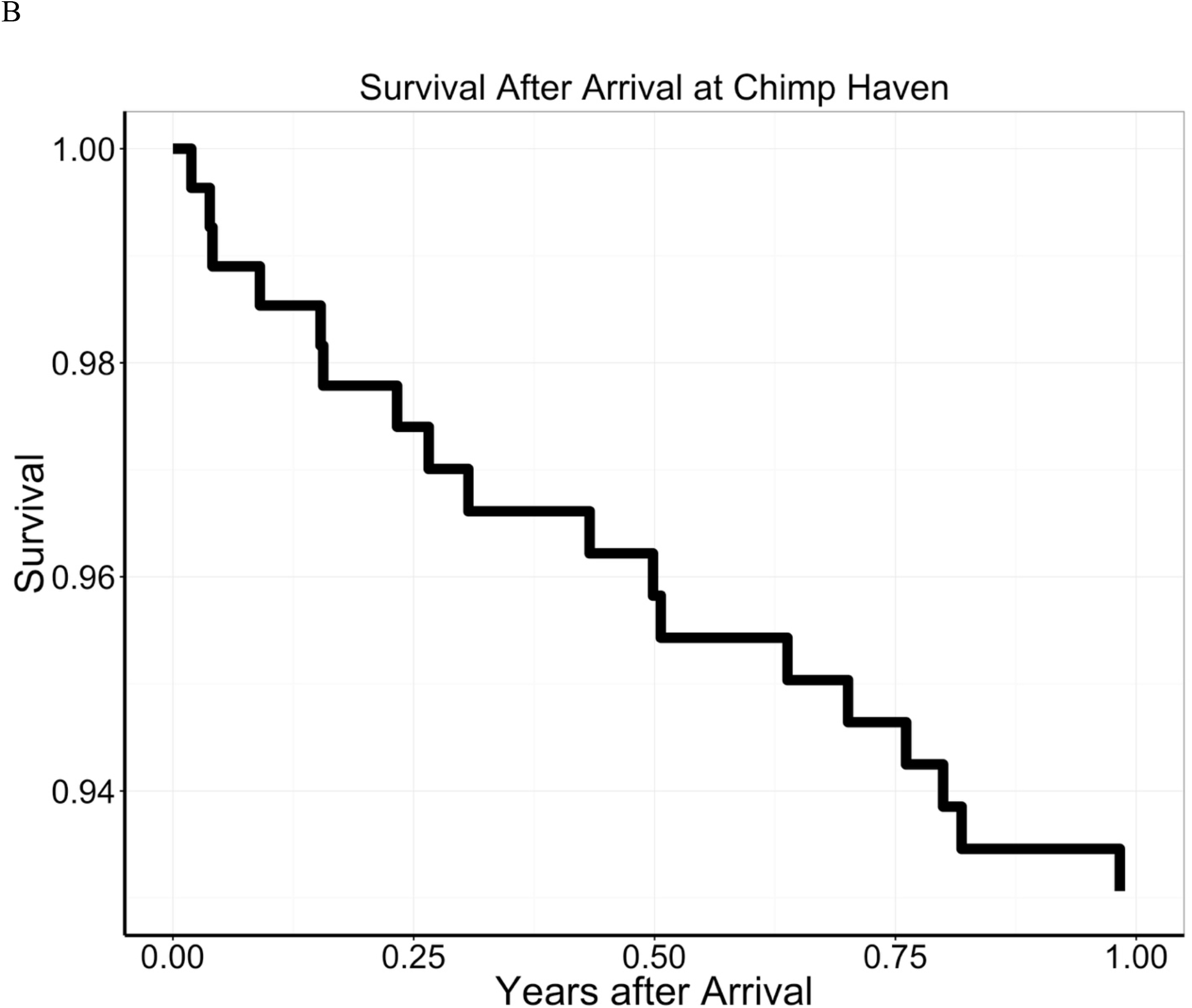
Kaplan-Meier survival plots for 273 chimpanzees transferred to Chimp Haven. Panel A shows total follow-up, while Panel B focuses on the first year after arrival.

## Discussion

Among 764 NIH-owned captive chimpanzees, location of residence was at most a modest correlate of all-cause mortality. Older age was by far the strongest predictor of mortality followed by male sex. Among 273 chimpanzees transferred to the federal sanctuary at Chimp Haven, older age was again a strong predictor of mortality. Other predictors, including season of transfer, and high- and low-ambient temperatures on day of arrival, were not significantly associated with mortality. Hazard for mortality followed a standard monotonic function after arrival. Mortality early – that is within 30 days – after transfer to Chimp Haven was unusual, accounting for only 3 of 107 deaths (<3%); two of the 3 deaths occurred in animals over 40 years of age. These findings argue against a strong association between location and mortality, and also argue against the existence of a materially increased hazard of early mortality following transfer as it has been recently practiced.

The analyses were conducted to inform NIH’s plans to retire its surviving chimpanzees. Transferring chimpanzees to a retirement sanctuary is a complicated and nuanced process, a process that by current practice takes into account social and veterinary health factors. The GAO recently recommended that the NIH develop and communicate a “clear implementation plan to transfer the remaining chimpanzees.”^8^ The current analysis will help NIH move to implement its retirement plan: decisions regarding the order of movement will focus largely on age, along with social groupings which are known to be important to well-being.^9^ We will also be able to use our regression models to estimate expected mortality over time.

Our analyses have a number of limitations. Most important, the only individual-level data available was date of birth, sex, vital status (along with date of death when applicable), and dates of residence at 4 specific locations. We did not consider specific infections (e.g. HCV, HIV), prior experiments, body mass, known chronic illness, laboratory findings (e.g. renal function), and details on social networks. Despite these limitations, our analyses argue against any strong associations between location or transfer and premature mortality. We plan to use our regression models to predict long-term survival among surviving chimpanzees at the federal sanctuary and other facilities, and in that way foster robust long-term plans for future transfers of chimpanzees remaining at other facilities.

